# Hepatocyte-like cells die via steroid hormone and nuclear receptor E75-mediated apoptosis

**DOI:** 10.64898/2025.12.29.696923

**Authors:** Devika Radhakrishnan, Noah Landgraf, Luigi Zechini, Alessandro Scopelliti, Neha Agrawal

## Abstract

Metabolic organs must sustain physiological function while retaining the capacity for timely cell death. Systemic hormones play a key role in coordinating this balance, yet how they regulate cell death in vivo remains unclear. Here, we investigate hormone-regulated cell death in a metabolically specialised organ using *Drosophila* oenocytes, polyploid hepatocyte-like cells, as a tractable in vivo model. Using non-invasive longitudinal live imaging combined with oenocyte-specific genetic manipulation, we directly visualise larval oenocyte death during metamorphosis. We show that larval oenocyte loss is a dynamic, multistep process controlled by the steroid hormone ecdysone. We further identify the ecdysone-induced nuclear receptor E75 as a key regulator of the timing of cell death, as loss of E75 triggers premature oenocyte death. Oenocyte-specific manipulation of cell death pathways, together with live imaging using genetically encoded caspase reporters, provides direct evidence that larval oenocytes die by apoptosis. Together, this work defines how systemic hormonal signals regulate the timing of apoptosis in metabolically specialised polyploid cells and establishes oenocytes as a powerful in vivo system for studying cell death in metabolic organs.

## INTRODUCTION

Metabolic organs must function efficiently while retaining the ability to remove damaged or unnecessary cells. In metabolic organs such as the liver, which undergoes extensive remodelling and cell death during development and in physiological and pathological contexts, failure to maintain this balance contributes to developmental defects and disease^1–5^. Hepatocytes, the principal functional cells of the liver, are highly metabolically active and require tight control of cell death to maintain tissue integrity and systemic homeostasis^6–8^. Hormonal signals play key roles in regulating hepatocyte function^8–11^.

However, how hepatocyte death is coordinated in vivo in response to systemic hormonal and physiological cues remains unclear. Moreover, most studies focus on pathological conditions, making it difficult to determine how hormonal signals regulate hepatocyte death under normal physiological states. Progress in this area has been limited by the lack of in vivo experimental systems that allow hormonal signalling and cell death dynamics to be examined together in metabolically specialised tissues in an intact animal.

*Drosophila melanogaster* provides a powerful genetic system to study how systemic hormonal signals regulate organ remodelling and cell death in vivo^12–16^. Metamorphosis is accompanied by extensive degradation and remodelling of larval organs coordinated by pulses of the steroid hormone 20-hydroxyecdysone (20E), commonly referred to as ecdysone^12–14,16^. Studies across multiple organs have shown that ecdysone-dependent tissue death is tightly spatiotemporally regulated and can involve apoptosis, autophagy, or combinations of both, depending on tissue context^14,17,18^. Ecdysone signalling acts through the heterodimeric ecdysone receptor complex (EcR)/Ultraspiracle (USP) to initiate transcriptional programmes that control developmental transitions^12,13^. Among the early ecdysone-responsive genes is the nuclear receptor E75^19,20^. E75 has been shown to modulate hormone-dependent transcription by antagonising EcR-driven activation in a tissue-and context-dependent manner^21^. Notably, *Drosophila* E75 is homologous to mammalian nuclear receptors^22^ such as members of the PPAR (peroxisome proliferator-activated receptor) family^23,24^ and NR1D2 (REV-ERBβ)^22,25^, which are key regulators of liver metabolism and hepatocyte survival^26–28^.

Despite extensive work on hormonally controlled tissue death, fundamental questions remain unresolved. It is still unclear how a single systemic hormone generates diverse, tissue-specific cell death responses, how the timing of death is determined, and why distinct death pathways are engaged in different organs. These questions are particularly important for metabolically specialised cells, where survival and death must be tightly coordinated with metabolic function and systemic physiology. They are also relevant to polyploid cells which often have altered sensitivity to death signals and can persist in pathological contexts such as cancer^29^.

Here, we use *Drosophila* hepatocyte-like cells, known as oenocytes, as a tractable in vivo system to study hormone-regulated cell death in a metabolically specialised tissue. *Drosophila* has a separate set of larval and adult oenocytes^30–32^. Larval oenocytes are a discrete population of large, polyploid cells with essential hepatocyte-like features, including metabolic specialisation, abundant smooth endoplasmic reticulum and peroxisomes, and roles in lipid regulation, starvation-induced lipid accumulation (steatosis), and fatty acid and hydrocarbon metabolism^30–38^.

Although larval oenocytes die during pupal development^38^, the cellular and molecular mechanisms underlying this process have not been defined. We have developed an experimental framework for non-invasive longitudinal imaging that enables continuous in vivo observation of the real-time dynamics of larval oenocyte cell death under physiological conditions. Using long-term live imaging combined with cell-type-specific genetic manipulation, we have found that oenocytes die via apoptosis and that this process is tightly regulated by ecdysone signalling and the nuclear receptor E75. Together, these findings identify a hormone-controlled mechanism that regulates apoptosis in hepatocyte-like cells and establish oenocytes as a powerful system to dissect how systemic endocrine cues govern cell death in metabolically specialised polyploid cells.

## RESULTS

### Live imaging reveals the cellular dynamics of larval oenocyte death

To understand the mechanisms driving oenocyte remodelling and physiological death, it is first necessary to define when and how this process occurs. This information is essential for mechanistic analysis, as oenocyte death must be coordinated with the remodelling of other larval organs and with changing metabolic demands during development. Unlike several larval organs such as the midgut, salivary glands, and tracheal system, which are removed early in metamorphosis^14,18,39–42^, larval oenocytes persist during pupal development^38^.

To characterise larval oenocyte removal comprehensively, we established a non-invasive, real-time longitudinal imaging strategy that allows larval oenocytes to be followed continuously in vivo throughout pupal development without affecting animal viability. Oenocytes were specifically labelled using an oenocyte-specific GAL4 driver (*PromE-GAL4*)^43,44^ to mark cell membranes and nuclei. Two complementary imaging approaches were combined to capture oenocyte behaviour across developmental stages (Figure 1A, B). Whole-pupa imaging with the pupal case intact enabled continuous, non-invasive observation from the white prepupal stage 0 hours after puparium formation (APF) under physiological conditions (Figure 1A; Video S1). In parallel, imaging after pupal case removal following head eversion provided high-resolution views of oenocyte dynamics from approximately 12 hours APF (Figure 1B; Video S2). The large size of oenocytes facilitates reliable visualisation of cell morphology and cell death over time. Together, these approaches allowed uninterrupted tracking of oenocyte behaviour over more than four days of development.

**Figure 1:**
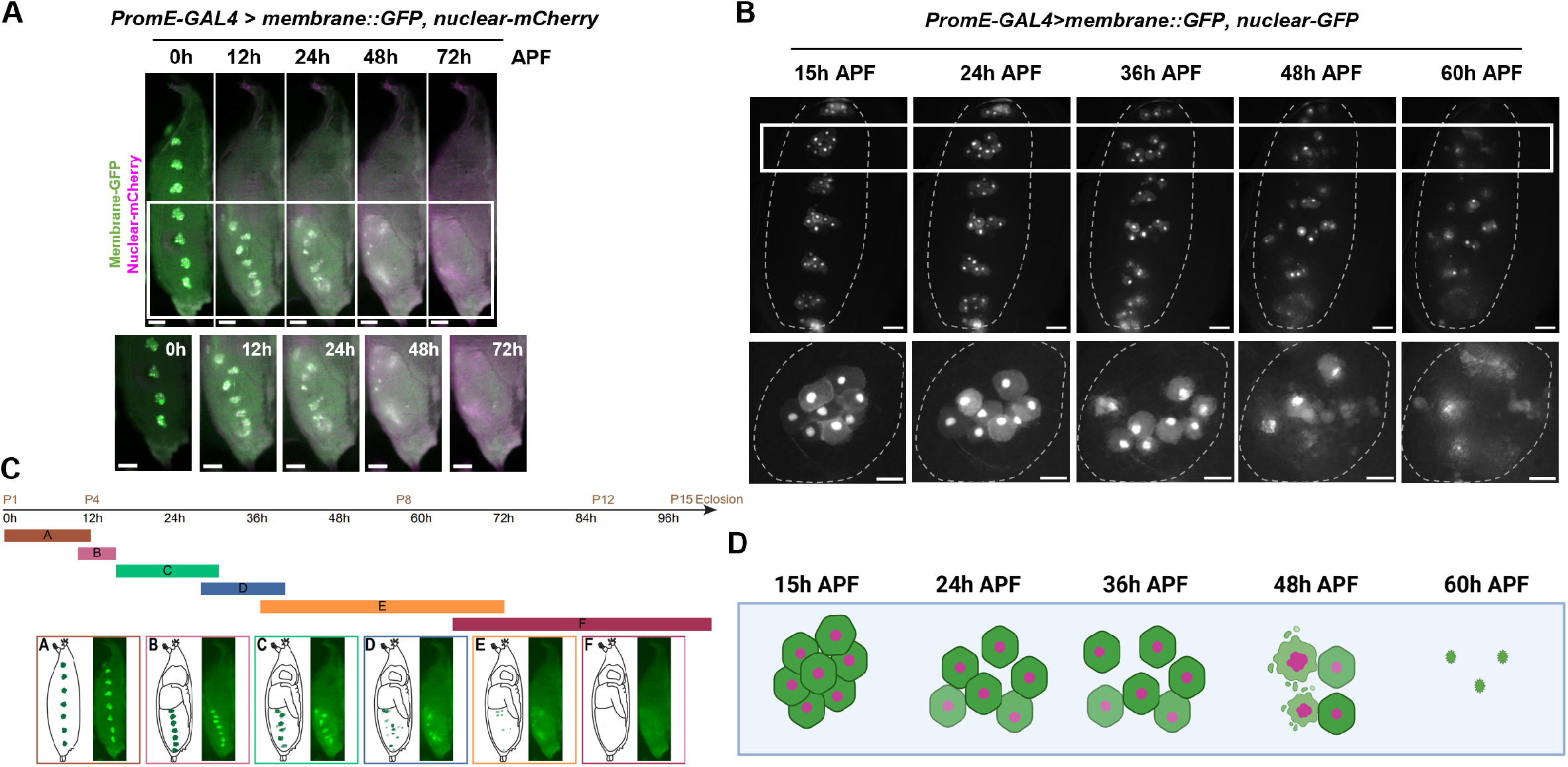
Live imaging reveals the cellular dynamics of larval oenocyte death during metamorphosis. **(A)** Top Panel: Still images from a time-lapse video (Video Sl) of larval oenocyte death in a pupa with intact pupal case. Images acquired every 20 minutes over a period of 72 hours. *PromE Ga/4>UASmCD8::GFP, UAS nuclear-mCherry* (membrane in green, nuclei in pink, and merge in white); APF: After Puparium Formation; Scale bar: 200 µm. Bottom Panel: Inset showing zoom-in of the pupal abdomen; Scale bar: 200 µm. **(B)** Top Panel: Still images from a time-lapse video (Video S2) of larval oenocyte cell death in a pupa imaged after pupal case removal. Images acquired every 20 minutes over a period of 72 hours. *PromE Ga/4>UASmCD8::GFP, UAS-nuclear-GFP* (membrane and nuclei in grayscale); Scale bar: 100 µm. Bottom Panel: Inset showing cellular dynamics of a single oenocyte cluster; Scale bar: 50 µm. **(C)** Timeline of cellular and developmental events underlying oenocyte loss during metamorphosis, divided into categories A-F, with representative pupae and schematics for each stage. **(D)** Schematic representation of the cellular dynamics of larval oenocyte death in a single larval oenocyte cluster over the course of metamorphosis. Created in Biorender.

Live imaging revealed that oenocyte degradation proceeds through a reproducible sequence of cellular and developmental events (Figure 1C, D). During the prepupal stage at 0 hours APF, oenocytes are organised into seven bilaterally symmetrical, segmentally repeated clusters of six to eight tightly associated cells positioned near the pupal surface (Category A). After head eversion, around 12 hours APF, these clusters reposition toward the developing abdomen (Category B). Individual oenocytes then progressively disengage from one another (which we refer to as oenocyte disassembly) around 24 hours APF, accompanied by clear changes in cluster organisation (Category C). Oenocytes subsequently move within the pupal abdomen (Category D), followed by fragmentation and death of oenocytes around 48 hours APF (Category E). By approximately 72 hours APF, larval oenocytes are no longer detectable (Category F).

The persistence of larval oenocytes into mid-pupal development provides an extended window for live imaging under physiological conditions. Together, these data establish larval oenocytes as an in vivo system to study the cellular dynamics and underlying mechanism of cell death in a metabolically active organ.

### Ecdysone signalling is required for larval oenocyte death

To dissect the mechanisms controlling oenocyte remodelling and physiological death, we used the oenocyte-specific *PromE-GAL4*. Careful examination of GFP reporter expression revealed that the *PromE-GAL4* is not active in developing adult oenocytes during pupal stages (data not shown), ensuring that all genetic manipulations were restricted to larval oenocytes. This approach enables precise genetic perturbation in fewer than 100 larval oenocyte cells within an otherwise intact animal, allowing cell-autonomous effects to be analysed. We followed oenocyte fate throughout pupal development using single time-point imaging at 24-hour intervals, complemented by long-term live imaging.

We next asked which signal initiates and drives the process of oenocyte loss. During metamorphosis, pulses of the steroid hormone ecdysone orchestrate larval organ remodelling and death, making ecdysteroid signalling a strong candidate regulator^12–14^. We therefore disrupted ecdysone signalling specifically within oenocytes by downregulating either the Ecdysone Receptor (*EcR*) or its co-receptor Ultraspiracle (*USP*) using *PromE-GAL4*. Loss of either *EcR* or *USP* strongly blocked oenocyte removal (Figure 2A, B). In both conditions, oenocytes persisted throughout pupal development, and pupae completed metamorphosis to form adults that surprisingly still retained larval oenocytes, as shown for *EcR* knockdown (Figure 2C).

**Figure 2:**
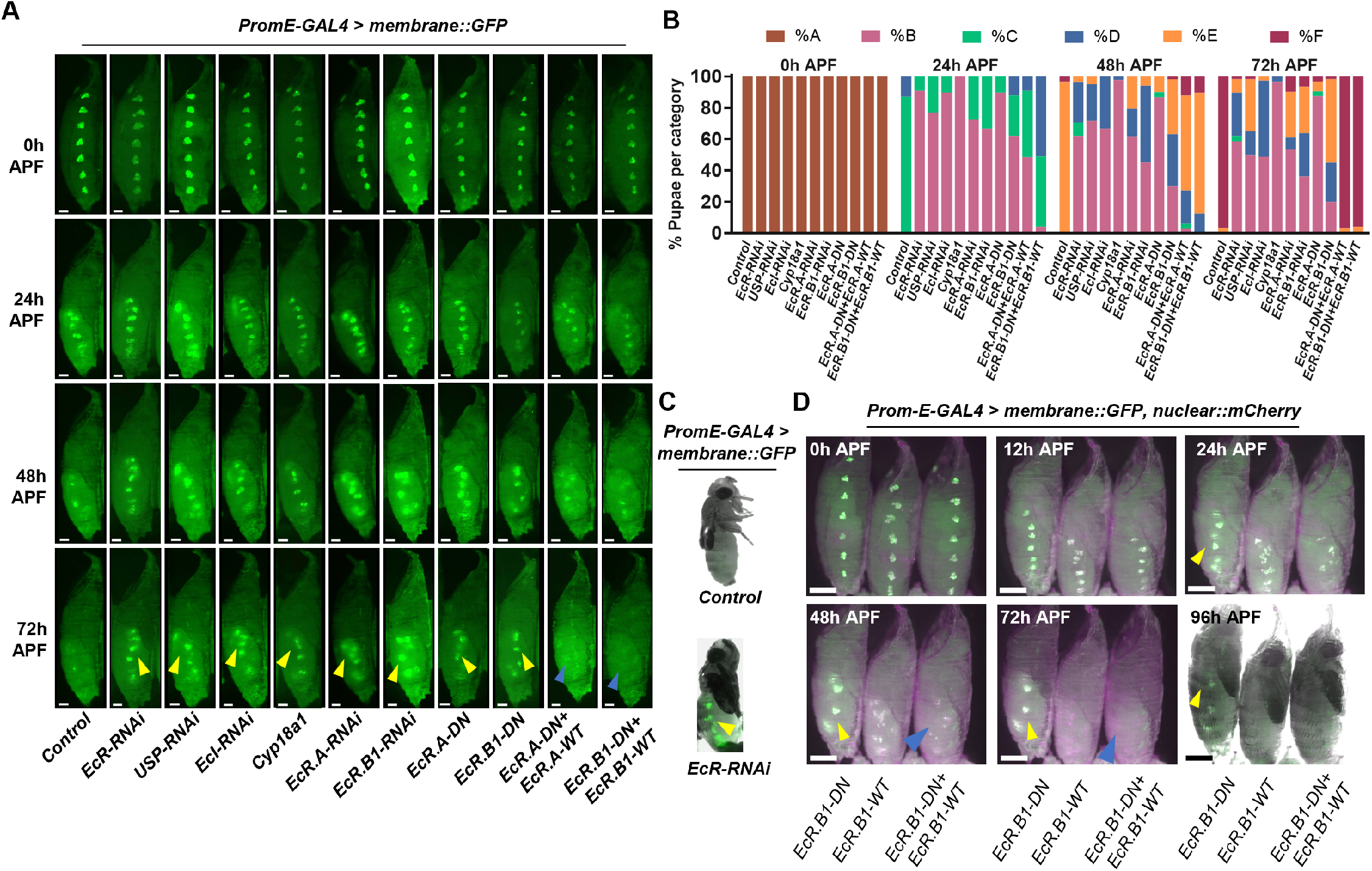
Ecdysone signalling regulates larval oenocyte cell death. (A) Representative images showing that oenocyte-specific downregulation of the Ecdysone Receptor *(EcR)*, Ultraspiracle *(USP)*, or Ecdysone Importer (fc/), as well as overexpression of *Cyp18a1*, prevents oenocyte death (yellow arrowheads), with larval oenocytes still remaining at 72hrs APF. Similarly, oenocyte-specific downregulation of *EcR*.*A* or *EcR*.*81* isoforms, or expression of a dominant-negative *EcR*.*A* or *EcR*.*81* transgene, blocks oenocyte removal (yellow arrowheads). Co-expression of the corresponding wildtype *EcR* isoforms in the dominant-negative background restores normal oenocyte death (blue arrowheads); Scale bar: 200 µm. (B) Percentage of pupae in each oenocyte degradation category at 0, 24, 48, and 72h APF for the indicated genotypes. *PromE-GAL4>UAS-mCD8::GFP* was crossed to *mCherryRNAi* (Control) *n* = 60, *EcR-RNAi n* = 58, *USP-RNAi n* = 60, *Ec/-RNAi n* = 39, *UAS-Cyp18a1 n* = 30, *EcR*.*A-RNAi n* = 53, *EcR*.*81-RNAi n* = 47, *EcR*.*A-DN n* = 32, *EcR*.*81-DN n* = 60, *EcR*.*A-DN; EcR*.*A-WT n* = 31, *EcR*.*81-DN; EcR*.*81-WT n* = 49. (C) Eclosed adult fly showing persistent larval oenocytes after oenocyte-specific EcR knockdown (yellow arrowhead), compared with control. (D) Still images from a time-lapse video (Video S3) of larval oenocyte cell death in pupae with intact pupal case of the indicated genotypes. Images acquired every 20 minutes over a period of 94 hours. Oenocyte death is blocked (yellow arrowhead) when *EcR*.*B1-DN* is expressed. Co-expression of the *EcR*.*B1-WT* in the dominant-negative background restores normal oenocyte removal (blue arrowheads). *(PromE Ga/4>UASmCD8::GFP, UAS-nuclear-mCherry* was crossed to *EcR*.*B1-DN* (left), *EcR*.*B1-WT* (middle), and *EcR*.*B1-DN; EcR*.*B1-WT* (right); membrane in green, nuclei in pink, and merge in white; Scale bar: 200 µm).

These results establish that ecdysone signalling acts cell-autonomously within oenocytes to drive their removal, extending the role of ecdysone-dependent tissue death to a metabolically specialised cell population that persists into mid-pupal development.

### Intracellular ecdysone availability modulates oenocyte death

A requirement for the EcR/USP receptor complex indicated that ecdysone signalling acts directly within oenocytes to regulate their death. This prompted us to ask whether intracellular ecdysone availability itself influences the progression of oenocyte loss. If ecdysone functions locally within oenocytes, then altering the amount of hormone available inside these cells should modify their ability to undergo death.

Intracellular ecdysone levels are regulated by multiple mechanisms, including hormone uptake and inactivation. The ecdysone importer EcI facilitates cellular uptake of ecdysone^45^, while the cytochrome P450 enzyme Cyp18a1 catalyses ecdysone inactivation^46^. Manipulating these components provides a means to alter effective hormone signalling within a specific cell type without broadly perturbing systemic endocrine cues. We therefore targeted these components specifically in oenocytes. Oenocyte-specific knockdown of *EcI* or expression of *Cyp18a1* strongly impaired oenocyte disassembly and death (Figure 2A, B). In both cases, oenocytes persisted with phenotypes closely resembling those observed following disruption of the EcR/USP receptor complex (Figure 2A, B).

Together, these findings show that oenocyte removal is highly sensitive to intracellular ecdysone levels and support a model in which direct, cell-autonomous ecdysone signalling regulates the timing and execution of oenocyte death during metamorphosis. The ability to follow oenocyte fate across mid-pupal development further allows changes in hormone availability to be directly linked to the dynamics of cell death in vivo.

### EcR.A and EcR.B1 isoforms are required for oenocyte death

Determining which EcR isoforms regulate oenocyte death is essential for understanding how a single systemic hormone generates tissue-specific outcomes during metamorphosis. In many larval tissues, the EcR isoform EcR.B1 is associated with death competence, whereas EcR.A has more often been linked to tissue remodelling or survival^12–14^. As oenocytes persist into mid-pupal stages, it is unclear which EcR isoform mediates this process.

To address this, we selectively disrupted *EcR*.*A* or *EcR*.*B1* within oenocytes and assessed oenocyte fate. Knockdown of either isoform blocked oenocyte loss (Figure 2A, B). Consistent with this, expression of dominant-negative forms^47^ of *EcR*.*A* or *EcR*.*B1* also prevented oenocyte removal, indicating that both isoforms contribute to this process (Figure 2A, B).

We confirmed the specificity of these effects using isoform-specific rescue experiments. Co-expression of wild-type *EcR*.*A* with *EcR*.*A-DN*, or wild-type *EcR*.*B1* with *EcR*.*B1-DN*, restored oenocyte removal in each case (Figure 2A, B), demonstrating that the dominant-negative phenotypes reflect isoform-specific loss of EcR function. Time-lapse live imaging further supported these conclusions. Oenocytes persisted in pupae expressing dominant-negative EcR isoforms, whereas they were efficiently removed in rescue conditions (Figure 2D; Video S3 shows *EcR*.*B1-DN, EcR*.*B1-WT*, and rescue).

Together, these data show that ecdysone signalling through both EcR.A and EcR.B1 are required for larval oenocyte death. This requirement indicates that oenocyte death does not follow the prevailing view that EcR.B1 alone confers death competence during metamorphosis.

### Oenocyte-specific downregulation of E75 accelerates oenocyte death

We next asked which downstream transcriptional regulators regulate oenocyte cell death. Broad-Complex (Br-C), E74A, and the nuclear receptor E75 are early ecdysone-induced factors that shape tissue-specific responses to ecdysone signalling^48^. Oenocyte-specific knockdown of *Br-C* or *E74A* had no detectable effect (Figure 3A, B). In contrast, *E75* knockdown tested with two independent RNAi lines caused a striking acceleration of oenocyte elimination. By 24 h APF, *E75*-depleted oenocytes had already fragmented and died, while control oenocytes remained largely intact (Figure 3A, B). In contrast, oenocyte-specific overexpression of *E75* delayed oenocyte removal (Figure 3A, B).

**Figure 3:**
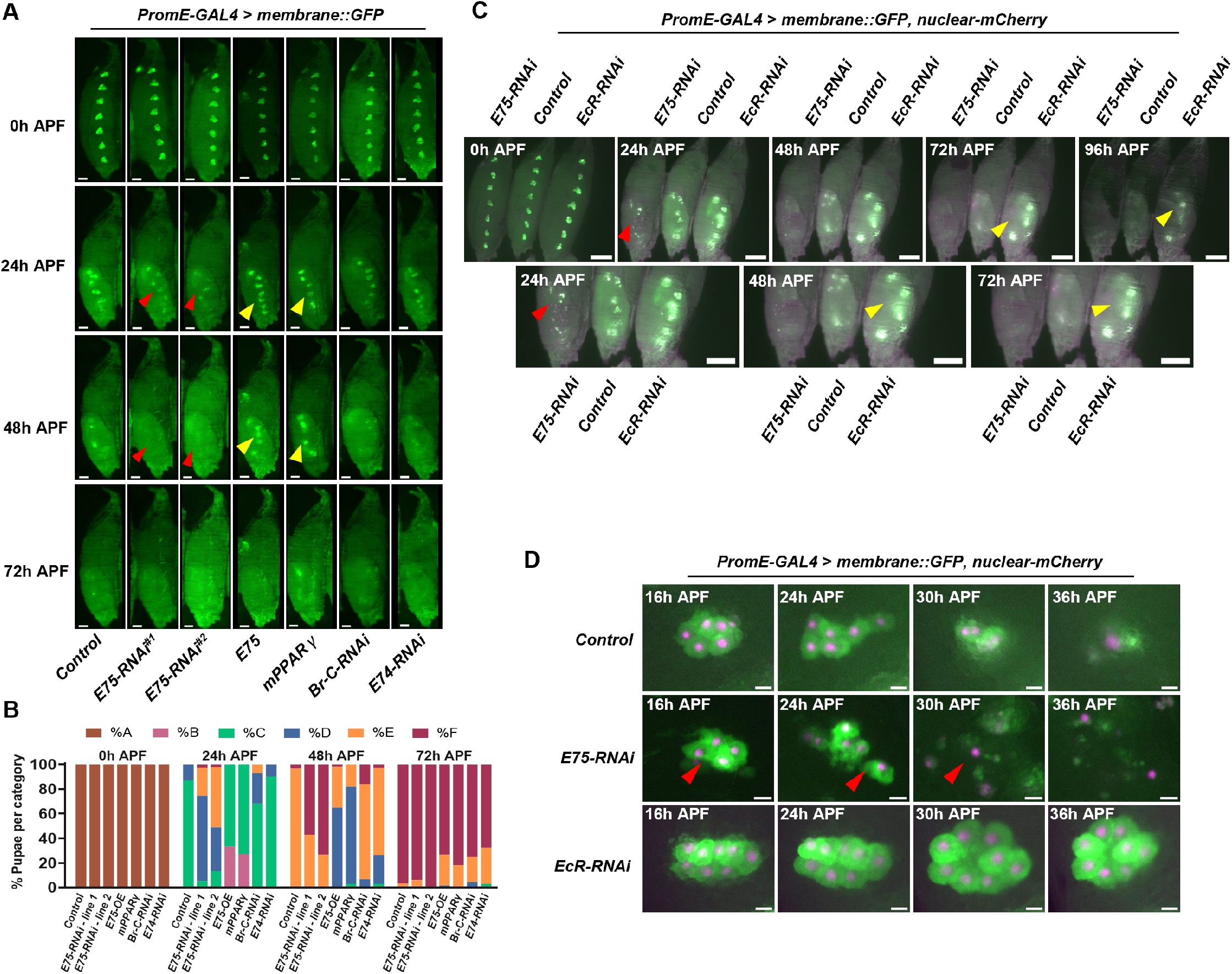
Oenocyte-specific *E75* knockdown accelerates oenocyte death. **(A)** Representative images of pupae with oenocyte-specific downregulation of nuclear receptor *E75* (with two independent RNAi lines) leads to accelerated cell death with oenocytes dying much more rapidly than in controls (red arrowheads). Overexpression of E75 or mouse peroxisome proliferator-activated receptor gamma *(mPPARy)* results in delayed oenocyte loss (yellow arrowheads). Downregulation of *BR-C* and *E74A* did not impact oenocyte removal. (Scale bar: 200 µm). **(B)** Percentage of pupae in each oenocyte degradation category at 0, 24, 48, and 72h APF for the indicated genotypes.: *PromE-GAL4>UAS-mCD8::GFP* was crossed to *mCherryRNAi* (Control) *n* = 60, *E75-RNAi (line 1) n =* 98, *E75-RNAi (line 2) n* = 45, *UAS-E75 n* = 60, *UAS-mPPARy n* = 33, *Br-C-RNAi n* = 44, *E74-RNAi n* = 34. **(C)** Still images from a time-lapse video (Video S4) of larval oenocyte cell death in pupae with intact pupal case of the indicated genotypes. Images acquired every 20 minutes over a period of 96 hours. Oenocyte death is blocked upon *EcR* knockdown (yellow arrowhead) but accelerated upon *E75* knockdown (red arrowhead). *(PromE Ga/4>UASmCD8::GFP, UAS-nuclear-mCherry* was crossed to *E75-RNAi* (left), *w1118* (control in the middle), *EcR-RNAi* (right); Membrane in green, nuclei in pink, and merge in white; Scale bar: 250 µm). Bottom panel: Inset showing zoom-in of the pupal abdomen with oenocyte specific *E75* knockdown and *EcR* knockdown at 24, 48, and 72h APF (Scale bar: 200 µm). **(D)** Single oenocyte cluster with oenocyte-specific *E75* or *EcR* knockdown at the indicated time points, following removal of the pupal case. Red arrowheads indicate rapid oenocyte loss following *E75* knockdown. *(PromE Ga/4>UASmCD8::GFP, UAS-nuclear-mCherry* was crossed to *w1118* (control on the top), *E75-RNAi* (middle), and *EcR-RNAi* (bottom); membrane in green, nuclei in pink, and merge in white; Scale bar: 25 µm).

E75 is homologous to mammalian nuclear receptors such as PPAR (peroxisome proliferator-activated receptor) family members^22–24^ which play key roles in liver and adipose metabolism. We next tested how overexpression of mouse PPARγ within oenocytes impacts oenocyte loss. Intriguingly, mouse PPARγ overexpression delays oenocytes removal (Figure 3A, B), suggesting functional conservation between E75 and PPARγ.

Whole-pupa live imaging with oenocyte-specific knockdown of *E75* or *EcR* together with control pupae allowed direct comparison of oenocyte behaviour under identical conditions, making differences in normal, accelerated, or blocked oenocyte death readily apparent (Figure 3C; Video S4). In pupae with oenocyte-specific *EcR* knockdown, oenocytes remained tightly clustered and failed to initiate removal. In contrast, *E75*-depleted oenocytes began fragmenting immediately after head eversion, and elimination was complete by ∼30 h APF. Continuous live imaging enabled direct visualisation of these distinct dynamic behaviours that would be difficult to capture using static time-point analysis alone. Imaging following pupal case removal confirmed rapid and extensive oenocyte fragmentation after *E75* knockdown (Figure 3D), compared with controls. However, oenocytes remained organised in stable clusters after *EcR* knockdown (Figure 3D).

We therefore find that *E75* knockdown results in premature oenocyte loss compared with controls, while overexpression of E75 or its mammalian homolog PPARγ delays oenocyte loss. Together, these data identify E75 as a key conserved regulator of oenocyte death.

### Apoptosis mediates larval oenocyte death during metamorphosis

Larval organ death during metamorphosis can proceed through apoptosis, autophagy-dependent cell death, or combinations of both, depending on tissue context^17,18^. Defining the cell death mechanism used by oenocytes is therefore necessary to understand how their death is hormonally regulated.

To distinguish between these possibilities, we first selectively disrupted core components of the autophagy pathway^49^ specifically in oenocytes. Knockdown of core genes of the autophagy pathway, including *Atg1, Atg7*, or *Atg8a* or combined knockdown of *Atg1* and *Atg8a*^50^ had no significant impact on oenocyte loss (Figure S1). In contrast, inhibiting apoptosis disrupted oenocyte death. Expression of the apoptosis inhibitors^51^ *Diap1*, microRNA that simultaneously inhibits *reaper, hid* and *grim* (miRHG)^52^, or *p35*, as well as knockdown of the initiator caspase *Dronc*, delayed or blocked oenocyte removal (Figure 4A, B).

**Figure 4:**
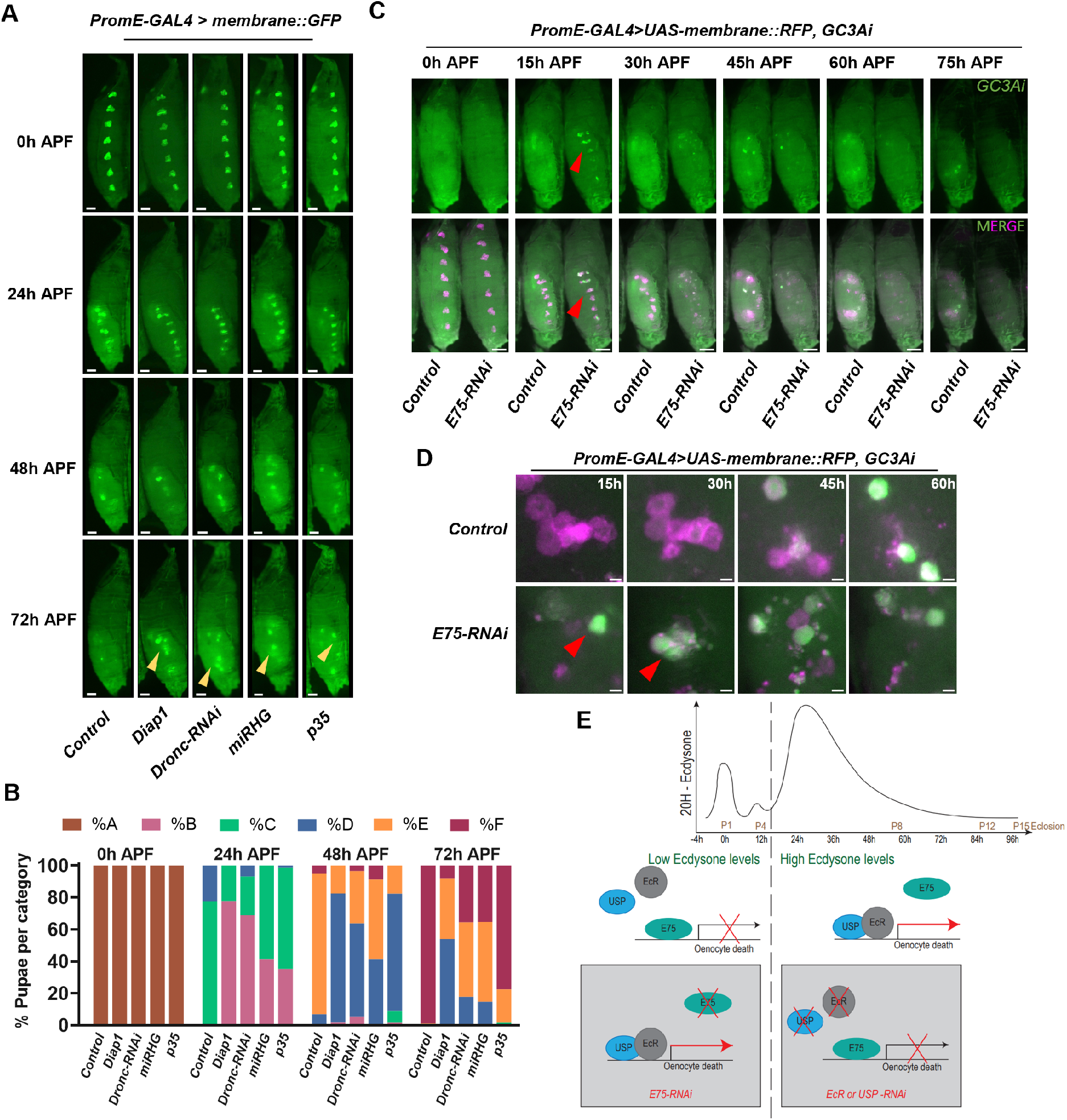
Apoptosis mediates larval oenocyte death during metamorphosis. **(A)** Representative images of pupae with oenocyte-specific inhibition of apoptosis. *Drone* knockdown or overexpression of *Diapl, p35* or microRNA targeting *reaper, hid* and *grim* [miRHG] delay or block oenocyte removal (Scale bar: 200 µm). **(B)** Percentage of pupae in each oenocyte degradation category at 0, 24, 48, and 72h APF for the indicated genotypes. *PromE-GAL4>UAS-mCD8::GFP* was crossed to *mCherryNLS* (Control) *n* = 58, *UAS-Diapl n* = 63, *Dronc-RNAi n* = 45, *UAS-miRHG n* = 34, *UAS-p35 n* = 67. **(C)** Still images from a time-lapse video (Video S5) of larval oenocytes expressing caspase reporter GC3Ai. Images acquired every 20 minutes over 90 hours. The pupa with *E75* knockdown shows GC3Ai expression soon after head eversion (red arrowhead) unlike controls. *(PromEGa/4>UASmCD8::RFP; UASGC3Ai* was crossed to *w1118* (control on the left), and *E75-RNAi* (right); Caspase reporter GC3Ai in green, membrane in pink, and merge in white; Scale bar: 250 µm). **(D)** Single oenocyte cluster (membrane-labelled with mRFP, magenta) expressing the caspase reporter GC3Ai (green), shown for control and oenocyte-specific *E75* knockdown at indicated time points. *E75* knockdown results in earlier GC3Ai expression (red arrowheads). *(PromE-Ga/4>UASmCD8::RFP; UASGC3Ai* was crossed to *w1118* (control on the top), and *E75-RNAi* (bottom); Caspase reporter GC3Ai in green, membrane in pink, and merge in white; Scale bar: 25 µm). **(E)** Working model for the role of *EcR/USP* and *E75* in the regulation of oenocyte death. The grey boxes indicate specific conditions leading to accelerated or blocked oenocyte death in response to knockdown of *E75* or *EcR/USP*, respectively (from left to right). We propose a model in which oenocyte death is controlled by the balance between EcR/USP-mediated activation and E75-dependent repression. During metamorphosis, smaller early ecdysone pulses are insufficient to overcome E75-mediated repression, allowing oenocytes to persist. The larger mid-pupal ecdysone peak shifts this balance toward EcR/USP activity, triggering oenocyte apoptosis. Consistent with this model, disruption of EcR/USP or specific EcR isoforms blocks oenocyte death, whereas loss of E75 removes this repression and permits premature oenocyte death at lower hormone levels.

These results demonstrate that larval oenocyte death proceeds primarily through apoptosis. This places oenocytes alongside dorsal external oblique muscles, which also undergo apoptosis during metamorphosis^40^, and contrasts with organs such as the salivary gland, midgut, and tracheal system, where autophagy contributes substantially to tissue histolysis^18,49^.

### Live caspase reporter imaging reveals oenocyte apoptosis

To understand the dynamics of oenocyte apoptosis, we next asked how apoptotic execution is initiated and progresses in vivo. We leveraged the oenocyte live-imaging system developed in this study to directly visualise caspase activation with high temporal resolution using the GFP-based caspase-3–like activity indicator (GC3Ai), a genetically encoded reporter that enables real-time detection of effector caspase activity in vivo without fixation^53^.

Interestingly, unlike controls, oenocyte-specific knockdown of *E75* triggered robust GC3Ai activation shortly after head eversion, coinciding with the accelerated oenocyte fragmentation observed in these animals (Figure 4C, D; Video S5). Together, these data identify E75 as a crucial determinant of the timing and mediator of oenocyte death. The oenocyte system is particularly powerful for revealing these dynamics, as genetically encoded caspase reporters allow apoptotic execution to be followed directly in individual cells within living animals.

## DISCUSSION

Our study shows that apoptotic cell death in a metabolically specialised tissue is a tightly regulated process that depends on systemic ecdysone signalling and the ecdysone-induced nuclear receptor E75. Our data support a working model in which oenocyte death is regulated by the balance between EcR/USP-mediated transcriptional activation and E75-mediated repression (Figure 4E). During metamorphosis, circulating ecdysone rises in a series of pulses, with two smaller peaks preceding a larger mid-pupal surge^12,13,54^. We hypothesize that during early pupal stages, E75-mediated repression of oenocyte death cannot be overcome by the smaller ecdysone peaks. Only when the higher mid-pupal ecdysone peak is reached does EcR/USP activity overcome E75-dependent repression, triggering oenocyte apoptosis. This model is strongly supported by our functional data: disrupting *EcR/USP* or specific EcR isoforms blocks oenocyte death, reflecting the necessity of ecdysone signalling for the activation of this process. Similarly, oenocyte-specific overexpression of *E75* delayed oenocyte removal. In contrast, *E75* knockdown removes its repressive control, allowing lower levels of ecdysone to prematurely trigger oenocyte loss, explaining the accelerated death observed.

This mechanism is consistent with earlier work in the larval salivary gland, where EcR/USP and E75 compete for shared regulatory regions and drive opposing transcriptional outcomes^21^. Similarly, loss of *E75* increases apoptosis in *Drosophila* tumour models^25^. In addition, heterochronic phenotypes in *E75* mutants^20^ and premature fat body dissociation following *E75* knockdown^55^ further support a broader, conserved role for E75 in controlling the timing of ecdysone-responsive developmental events.

While the salivary glands, midgut, and tracheal system are removed early in metamorphosis^18^, we confirm that larval oenocytes persist into mid-pupal development. This delayed removal likely reflects their role in the production of cuticular hydrocarbons^34^. We further hypothesize that prolonged oenocyte survival may contribute to fat body remodelling ^56^ by regulating lipid metabolism within the metabolically closed environment of the pupa^57^.

Together, these findings highlight a major strength of the oenocyte system. As oenocytes persist into mid-pupal stages, apoptotic execution can be followed continuously in living animals using real-time reporters. Many larval tissues are eliminated earlier^14,18^, when live imaging is technically limited, restricting analysis of apoptotic dynamics. Oenocytes therefore provide a particularly accessible system for studying how systemic endocrine cues are translated into precisely timed cell death decisions in vivo.

Finally, our findings extend beyond *Drosophila* metamorphosis. E75 is homologous to mammalian nuclear receptors such as members of the PPAR family^23,24^, key regulators of lipid metabolism and liver physiology^26–28^. Our observation that oenocyte-specific overexpression of mammalian PPARγ can significantly delay oenocyte loss supports functional conservation between E75 and PPARγ and shows that a mammalian E75 ortholog can alter the apoptotic programme in *Drosophila* oenocytes.

Oenocytes are also a tractable model for studying mechanisms that determine the persistence or death of polyploid cells, a cellular state frequently associated with therapy resistance in cancer^58^. Moreover, they also provide a developmental model for metabolically specialised post-mitotic cells that play a transient role in mammalian development prior to cell death. Relevant mammalian examples include foetal adrenal zone cells, which perform lipid -metabolising endocrine functions during development and undergo apoptosis after birth^59^, and trophoblast giant cells, which are large, polyploid, placental cells with specialised metabolic functions^60^. The oenocyte system thus offers a powerful platform to explore how hormonal, metabolic, and transcriptional signals are integrated to control cell survival and death in vivo in metabolically specialised polyploid cells.

## METHODS

### Fly husbandry and fly stocks

Fly stocks were reared on a modified *Drosophila* Lewis medium^61^. All experimental flies were kept in incubators at 25°C on a 12-hour light/dark cycle. Unless otherwise stated, all stocks were obtained from the Bloomington *Drosophila* Stock Centre. Bloomington stock numbers are provided in brackets.

*PromE-Gal4* (65404, 65405), *UAS-mCD8::GFP* (5137), *UAS-mCherry-NLS* (38424), *UAS-mCD8-mRFP, UAS-mCherry-RNAi* (35785), *UAS-EcR-RNAi* (9327, 9326), *UAS-EcI-RNAi*^45^ *and UAS-Cyp18a1*^46^ (gift from Naoki Yamanaka), *UAS-EcR*.*A*.*RNAi* (9328), *UAS-EcR*.*B1*.*RNAi* (9329), *UAS-EcR*.*A*.*DN* (9452), *UAS-EcR*.*B1-DN* (6869), *UAS-EcR*.*A-WT* (6470), *UAS-EcR*.*B1-W*T (6469), *UAS-E75-RNAi –* line 1 (26717), *UAS-E75-RNAi* – line 2 (VDRC – 44851), *UAS-E75::FLAG* (gift from Tobias Reiff, original source^62^), *UAS-mPPAR*γ (83644), *UAS-Br-C-RNAi* (51189), *UAS-E74-RNAi* (29353), *UAS-Diap1* (6657), *UAS-Dronc-RNAi* (32963), *UAS-miRHG*^52^, *UAS-p35* (5072), *UAS-GC3Ai* (84343), *UAS-Atg1-RNAi* (44034), *UAS-Atg7-RNAi* (27707), *UAS-Atg8a-RNAi* (28989).

### In vivo live imaging

Prepupae were gently collected with a wet paintbrush, rinsed in water, and dried. For imaging with the pupal case intact, pre-pupae were placed laterally on small droplets of halocarbon oil (Merck Life Science Limited, H8873) on a glass-bottom dish, ensuring the spiracles remained free of oil. For imaging with the pupal case removed, prepupae were dried overnight, placed ventral side down on double-sided tape, and the case was carefully removed using forceps. Pupae were then mounted laterally on a thin strip of Elmer’s glue^63^ applied to a petri dish lid, with the head positioned away from the surface. In both preparations, a damp tissue was included in the sealed dish to maintain humidity during imaging at 25°C.

### Time-point imaging and analysis

Pupae of the correct genotype were collected within a 6-hour window for 0 h APF imaging and attached laterally to a plastic petri dish lid using Elmer’s glue. A damp tissue was placed inside the dish to maintain humidity, and samples were kept at 25°C. The same pupae were imaged every 24 hours at the midpoint of the collection window up to 72 h APF, using identical imaging settings across genotypes and time points. Although most pupae completed metamorphosis, minor pupal lethality was observed. For quantification of oenocyte degradation, only pupae that reached stage P8 (50–72 h APF)^64^, defined by visible eye pigmentation, were included.

### Imaging & Image Analysis

Imaging was conducted using a Leica M205 FCA fluorescence stereomicroscope. Images were captured at a single z-plane every 10 or 20 minutes, as indicated in the figure legends over 72–96 hours. Image analysis was carried out in Fiji, and graphs were generated using GraphPad Prism.

## ACKNOWLEDGEMENTS

We thank Nuria Romero, Naoki Yamanaka, Andrea Brand, Catherine Davidson, Pierre Léopold, Alex Gould, Tobias Reiff, Andrew Davidson, Dhanya Cheerambathur, and all members of the lab for insightful discussions and for generously sharing resources. We thank David Kelly and Toni McHugh in the Light Microscopy Core at the Wellcome Discovery Research Platform for Hidden Cell Biology for technical assistance. We are also grateful to Flybase, the Bloomington *Drosophila* Stock Centre, and Vienna *Drosophila* Research Centre.

D.R. is supported by a Darwin Trust of Edinburgh PhD Studentship. This work was supported by funding from the University of Edinburgh Start-up Grant, Wellcome Trust Institutional Strategic Support Fund (ISSF3) award (IS3-R1.24 22-23), Medical Research Scotland Early Career Research Grant (ECG-1775-2021), Royal Society Research Grant (RGS\R2\222285) and an Academy of Medical Sciences Springboard Award (SBF008\1166).

## AUTHOR CONTRIBUTIONS

Conceptualization: D.R., N.L. and N.A.; Methodology: D.R., N.L. and N.A.; Validation: D.R. and N.L.; Formal Analysis: D.R., N.L. and N.A.; Investigation: D.R. and N.L.; Resources: D.R., N.L., L.Z., A.S. and N.A; Writing: D.R. and N.A.; Visualization: D.R., N.L. and N.A.; Supervision: N.A.; Project administration: N.A.; Funding acquisition: N.A.

## DECLARATION OF INTEREST

The authors declare no competing interests.

## SUPPLEMENTAL INFORMATION

**Figure S1:**
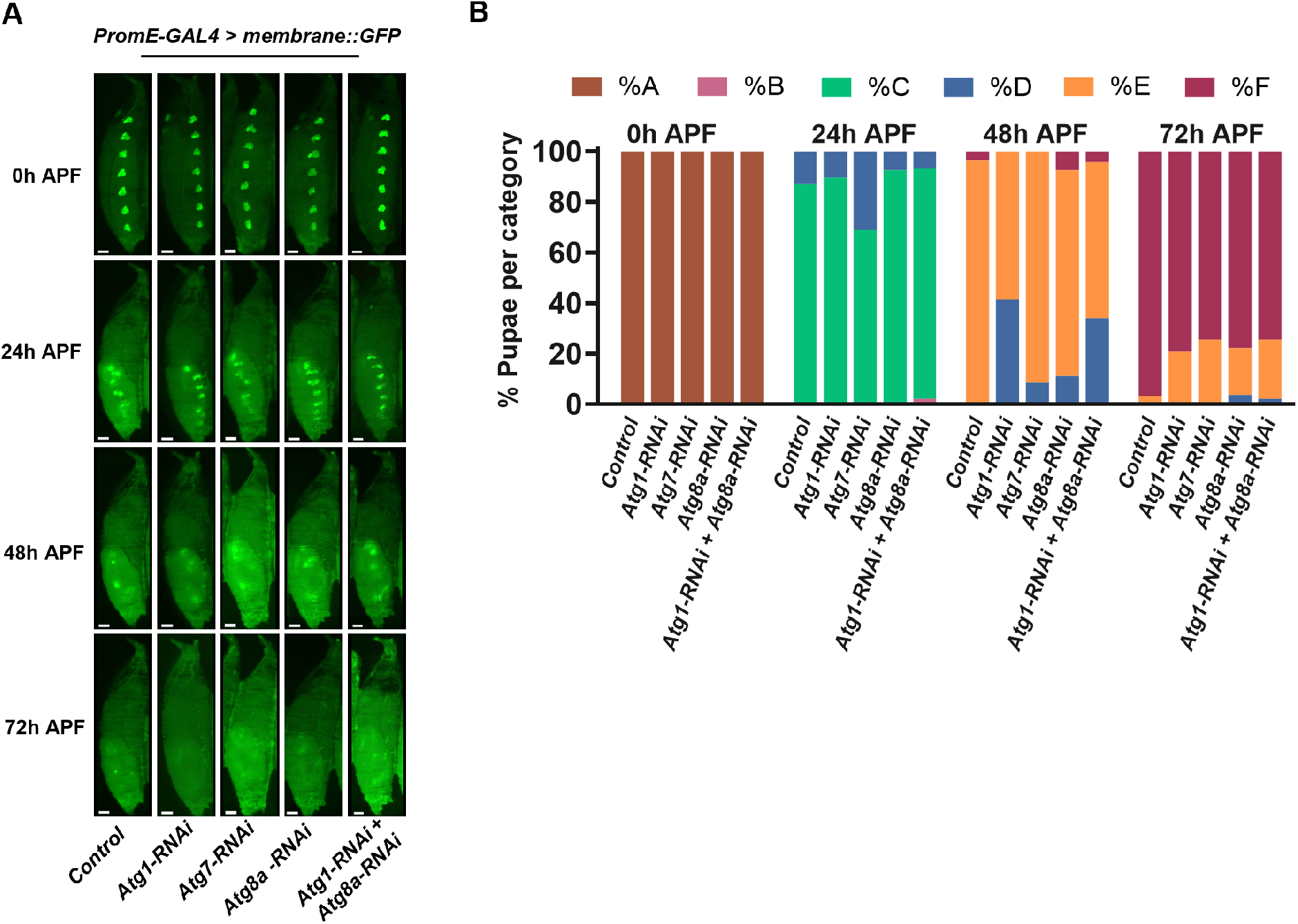
Oenocyte-specific downregulation of autophagy genes does not impact oenocyte death. **(A)** Representative images of pupae with oenocyte-specific downregulation of core autophagy pathway genes. (Scale bar: 200 µm). **(B)** Percentage of pupae in each oenocyte degradation category at 0, 24, 48, and 72h APF for the indicated genotypes. *PromE-GAL4>UAS-mCDB::GFP* was crossed to *mCherryRNAi* (Control) *n* = 40, *Atg1-RNAi n* = 29, *Atg7-RNAi n* = 47, *AtgBa-RNAi n* = 27, *Atg1-RNAi; AtgBa-RNAi n* = 47.

## SUPPLEMENTAL INFORMATION

**Video S1: Visualization of the cellular dynamics of larval oenocyte cell death in whole pupa**.

Time-lapse video of pupa with intact pupal case. Images acquired every 20 minutes over a period of 72 hours. *(PromE-Ga/4>UASmCD8::GFP, UAS-nuclear-mCherry* (membrane in green, nuclei in pink, and merge in white); Scale bar: 200 µm; Frame rate: 10 fps).

**Video S2: Visualization of the cellular dynamics of larval oenocyte cell death in pupa without pupal case**.

Time-lapse video of a pupa imaged after pupal case removal post head eversion. Images acquired every 20 minutes over a period of 72 hours. *(PromE-Ga/4>UASmCD8::GFP, UAS nuclearGFP* (shown in grayscale); Scale bar 100 µm; Frame rate: 10 fps).

**Video S3: Co-expression of wild-type *EcR*.*81* in a dominant-negative background restores oenocyte death**.

Time-lapse video of larval oenocyte cell death in pupae of indicated genotypes with intact pupal case. Images acquired every 20 minutes over a period of 94 hours. Oenocyte death is blocked when *EcR*.*81-DN* is expressed, and this phenotype is restored by co-expressing wild type *EcR*.*81* in a dominant negative background *(PromE-Ga/4>UASmCD8::GFP, UAS nuclear-mCherry* was crossed to *EcR*.*81-DN* (left), *EcR*.*81-WT* (middle), and *EcR*.*81-DN; EcR*.*81-WT* (right); membrane in green, nuclei in pink, and merge in white; Scale bar: 200 µm; Frame rate: 10 fps).

**Video S4: *E75* knockdown results in accelerated oenocyte death while oenocyte removal is blocked upon *EcR* knockdown**.

Time-lapse video of larval oenocyte cell death in pupae of indicated genotypes with intact pupal case. Images acquired every 20 minutes over a period of 96 hours. *E75*-depleted oenocytes begin fragmenting and dying immediately after head eversion (pupa on left) as compared to control (pupa in middle). In contrast, oenocytes remained tightly clustered and did not die upon oenocyte-specific *EcR* knockdown (pupa on right); *(PromE Ga/4>UASmCD8::GFP, UAS-nuclear-mCherry* was crossed to *E75-RNAi* (left), *w1118* (control in the middle), *EcR-RNAi* (right); membrane in green, nuclei in pink, and merge in white; Scale bar: 250 µm; Frame rate: 10 fps).

**Video S5: Oenocyte-specific *E75* knockdown leads to early induction of apoptosis**. Time-lapse video of larval oenocytes expressing caspase reporter GC3Ai (in green). Images acquired every 20 minutes over 90 hours. Unlike controls, *E75* knockdown in oenocytes induced robust GC3Ai activation shortly after head eversion, alongside accelerated oenocyte fragmentation. *(PromE-Ga/4>UASmCD8::RFP; UASGC3Ai* was crossed to *w1118* (control on the left), and *E75-RNAi* (right); Caspase reporter GC3Ai in green, membrane in pink, and merge in white; Scale bar: 250 µm; Frame rate: 10 fps).

## Notes

### Competing Interest Statement

The authors have declared no competing interest.

### Summary of Updates

New results added assessing the impact of E75, mammalian PPARγ, and miRHG (microRNA that simultaneously inhibits reaper, hid and grim) overexpression on oenocyte death. Figures and videos have been reorganised to improve clarity and flow, with additional detail added.

